# CLEM-Reg: An automated point cloud based registration algorithm for correlative light and volume electron microscopy

**DOI:** 10.1101/2023.05.11.540445

**Authors:** Daniel Krentzel, Matouš Elphick, Marie-Charlotte Domart, Christopher J. Peddie, Romain F. Laine, Cameron Shand, Ricardo Henriques, Lucy M. Collinson, Martin L. Jones

**Affiliations:** Imaging and Modeling Unit, Institut Pasteur, Université Paris Cité, Paris, France; Electron Microscopy Science Technology Platform, The Francis Crick Institute, London, UK; Cancer Dynamics Laboratory, The Francis Crick Institute, London, UK; UCL-Laboratory for Molecular Cell Biology, University College London, London, UK; Software Engineering & AI Science Technology Platform, The Francis Crick Institute, London, UK; Instituto de Tecnologia Química e Biológica António Xavier, Universidade Nova de Lisboa, Portugal

## Abstract

Correlative light and volume electron microscopy (vCLEM) is a powerful imaging technique that enables the visualisation of fluorescently labelled proteins within their ultrastructural context on a subcellular level. Currently, expert microscopists align vCLEM acquisitions using time-consuming and subjective manual methods. This paper presents CLEM-Reg, an algorithm that automates the 3D alignment of vCLEM datasets by leveraging probabilistic point cloud registration techniques. These point clouds are derived from segmentations of common structures in each modality, created by state-of-the-art open-source methods, with the option to leverage alternative tools from other plugins or platforms. CLEM-Reg drastically reduces the time required to register vCLEM datasets to a few minutes and achieves correlation of fluorescent signal to sub-micron target structures in EM on three newly acquired vCLEM benchmark datasets (fluorescence microscopy combined with FIB-SEM or SBF-SEM). CLEM-Reg was then used to automatically obtain vCLEM overlays to unambiguously identify TGN46-positive transport carriers involved in the trafficking of proteins between the trans-Golgi network and plasma membrane. The datasets are available in the EMPIAR and BioStudies public image archives for reuse in testing and developing multimodal registration algorithms by the wider community. A napari plugin integrating the algorithm is also provided to aid end-user adoption.

## Introduction

Correlative light and electron microscopy (CLEM) is a powerful imaging technique that seeks to capitalise on the advantages of light microscopy (LM) and electron microscopy (EM) while circumventing the drawbacks of each. This has made CLEM the imaging technique of choice to target rare and dynamic biological events that necessitate structural analysis at high resolution (1,2). Fluorescence microscopy (FM) is an LM imaging modality which generates contrast by tagging macromolecules in living cells and tissues with fluorescent proteins, enabling dynamic observation of their biological interactions. However, due to the diffraction limit of light, traditional FM cannot achieve a resolution better than around 200 nm, hindering fine structural details from being resolved (3). While super-resolution techniques can surpass this diffraction limit, such methods require specialised instruments, as well as specific sample preparation and imaging protocols, imposing additional constraints on the type of biological events that can be imaged (4). Moreover, FM generally tags specific macromolecules, therefore providing excellent molecular specificity. However, unlabelled structures cannot be observed. EM addresses these limitations, achieving orders of magnitude higher resolution while revealing the underlying biological context (5) in exquisite detail, but at the cost of a smaller field-of-view (FOV) and the lack of molecular specificity. By harnessing the complementary information derived from the correlation of LM and EM, CLEM has led to a variety of biological discoveries, such as establishing the structure of tunnelling nanotubes in neurons (6), observing blood vessel fusion events in zebrafish (2) and localising tuberculosis bacteria in primary human cells (1).

One of the most common approaches for performing CLEM experiments is to sequentially image a sample using FM and then EM, necessitating a two-stage sample preparation process. First, relevant structures are tagged with organic dyes or fluorescent proteins, and an image stack through the sample volume is acquired using FM. The sample is fixed with cross-linkers, either before or after the FM imaging, to conserve structural features. It is then stained with heavy-metal salts to introduce contrast, dehydrated, embedded in resin, and trimmed to the region of interest (ROI) (7). In volume EM (vEM), layers of the embedded sample are physically removed, and either the face of the block or the sections themselves are imaged to create a stack of serial EM images through the sample volume (5). This results in two corresponding image stacks, one from FM and one from EM, each containing complementary data from the same physical region of the sample, but typically imaged in different orientations. In addition to this orientation mismatch, the sample preparation and imaging processes can introduce both linear and non-linear deformations between the acquired FM and EM image volumes. Thus, to correlate the FM signal to the underlying structures visible in EM, the image volumes need to be brought into alignment. Due to the significant differences in resolution, contrast and FOV between FM and EM, this is a challenging task that cannot be directly approached with intensity-based methods that are routinely used for aligning data from visually similar modalities, e.g. magnetic resonance imaging (MRI) and computed tomography (CT).

There are two general approaches to solving this problem. The first approach is to process one or both images such that they share a similar visual appearance, for example, by directly converting across modalities (8–10) or by constructing a shared modality-agnostic representation (11,12). Once the processed image stacks are sufficiently similar in visual appearance, traditional intensity-based registration techniques, such as those employed in medical imaging (13), can be used to automatically align the two datasets. The second approach uses a landmark-based method, such as those implemented in software tools like BigWarp (14) and ec-CLEM (15). These tools rely on manually identifying precise spatial regions visible in both modalities, for example, small sub-cellular structures or prominent morphological features. By manually placing a landmark at an identical physical position in each modality, spatial correspondences can be established. A set of these landmarks can then be used to compute the transformation between the image volumes (1), bringing them into alignment via an iterative optimisation process (16,17). Methods of automating the selection of corresponding landmarks across modalities have been investigated, for example, determining cell centroids in LM and EM (18), or using semi-automated feature detection (e.g. AutoFinder in ec-CLEM) (15). However, such methods are often restricted to 2D or limited to relatively coarse alignment, requiring subsequent manual refinement. Moreover, deep learning-based methods that convert across modalities (8) or construct a shared modality-agnostic representation (12) rely on the availability of large amounts of aligned ground truth data. Due to the low throughput and required expertise of manual volume CLEM (vCLEM) alignment, generating such ground truth data is challenging.

A significant advantage of landmark-based approaches is the inherent sparsity of the representation, which substantially reduces memory and computational requirements compared to intensity-based registration techniques that must generally hold both image volumes in working memory. However, this manual landmark selection step is laborious and time-consuming, severely impacting throughput and potentially introducing bias, since the target structure is often directly used for registration. To avoid such biases, landmarks used for registration should be different from the target structures being studied wherever possible, taking care to ensure colour-correction between the channels in FM to avoid spectrally induced shifts in focal depth. Due to these limitations, robust and objective automation of landmark detection is highly desirable.

Here, CLEM-Reg is introduced, an automated vCLEM registration algorithm that relies on extracting landmarks from common structures in each modality. To achieve this aim, a workflow to segment mitochondria was developed. Mitochondria were chosen specifically because they are abundant and typically well-distributed across cells, and easily imaged in both FM and EM, enabling robust matching across modalities. Various segmentation approaches are routinely used in microscopy, ranging from classical image processing techniques (9,19,20) to machine learning (21–23). Depending on the complexity and density of structures in the image at hand, different algorithms are appropriate. For instance, segmentation of objects like mitochondria in FM acquisitions can be performed by filtering and thresholding coupled with further downstream image processing (19), while automatically segmenting EM data typically requires advanced deep learning techniques, due to the specific type of contrast obtained in EM imaging. Common deep learning architectures like “U-Net” (24) can be trained locally from scratch to very good effect, but the burden of obtaining sufficient ground truth data presents a huge challenge, often requiring significant amounts of expert effort or crowdsourcing of manual annotations (23). Recently, however, pre-trained “generalist” models such as MitoNet (25), based upon a “Panoptic-DeepLab” architecture (26), are able to provide out-of-the-box performance levels for mitochondrial segmentation in EM that are sufficient for many tasks, with the option to fine-tune on local data where necessary.

After segmentation, points are equidistantly sampled from the surface of the resulting mitochondria segmentations in both the FM and EM volumes, resulting in a “point cloud” for each modality. Point clouds are an attractive modality agnostic representation due to their inherent sparsity and the availability of a range of performant registration algorithms (16,27–29). However, unlike manually generated pairs of points, there is no guarantee of a one-to-one precise spatial correspondence between points in the different modalities. The coherent point drift (CPD) (27) algorithm overcomes this limitation by casting the alignment task as a probability density estimation problem, thereby removing the constraint of strict point correspondences.

Assessing registration performance in vCLEM overlays is challenging. Unlike in the medical imaging field where multimodal registration of MRI and CT scans is achieved by minimising mutual information (MI), no equivalent metrics exist to compare FM and EM volumes. This lack of metrics is an important hurdle in addressing automated vCLEM alignment and evaluating registration performance. Currently, the assessment of vCLEM overlay quality is based on qualitative inspection by experts, oftentimes by the same person who generated the alignment in the first place.

The aim of vCLEM experiments is to functionally label ultrastructure in EM with fluorescent signal. These structures can range from microns to a few nanometres in size. To test the limits of CLEM-Reg, registration performance was assessed on some of the smallest known organelles, namely lysosomes, which have a size of 0.3 to 1 µm (30,31). To quantify registration performance, two new metrics based on the correlation of fluorescent signal (Lysotracker) to sub-micron target structures (lysosomes) in EM are introduced here. Specifically, the volume of lysosomes overlaid by fluorescence was computed and centroid distances between fluorescent signal and lysosomes calculated. Since CLEM-Reg, to our knowledge, is the first algorithm to automate vCLEM alignment, manual registration by an expert was used as a baseline for comparison.

Performance quantification was conducted on two newly acquired vCLEM benchmark datasets using two different EM modalities (FIB-SEM and SBF-SEM). A third vCLEM dataset was acquired to investigate a rare and dynamic cellular process involving sub-micron organelles. TGN46 (TGN38 in rodents) has previously been observed in transport carriers involved in the trafficking of proteins between the trans-Golgi network (TGN) and plasma membrane (32–35). These transport carriers are rare, as they account for only 1 to 5% of total TGN46 signal (35). Here, CLEM-Reg is used to automatically register GFP-TGN46 signal to EM ultrastructure in 3D thereby facilitating accurate identification of TGN46 transport carriers that would have been missed by visual inspection of the EM volume alone. All datasets and overlays are made available to the community on EMPIAR and BioStudies (**Data and Code Availability**).

## Results

### Benchmark dataset acquisition

To assess the performance of CLEM-Reg against an expert, three benchmark vCLEM datasets (EMPIAR-10819, EMPIAR-11537 and EMPIAR-11666) of HeLa cells were acquired. Mitochondria (Mitotracker Deep Red), nucleus (Hoechst 33342), Golgi apparatus protein TGN46 (GFP-TGN46) and lysosomes (Lysotracker Red) in EMPIAR-10819 and EMPIAR-11666 and plasma membrane (WGA) in EMPIAR-11537 were tagged so that the unbiased registration performance could be assessed using target structures. After imaging the samples with a Zeiss Airyscan LSM900 microscope, two corresponding EM volumes (EMPIAR-10819 and EMPIAR-11537) with an isotropic voxel-size of 5 nm were acquired using a focused ion beam scanning electron microscope (FIB-SEM). The EM volume in EMPIAR-11666 was acquired using a serial block-face scanning electron microscope (SBF-SEM) with a voxel-size of 7 nm in XY and 50 nm in Z. The acquired images were pre-aligned following a routine image processing workflow in Fiji (**Materials and Methods**, **Fig. 1A**).

**Figure 1.**
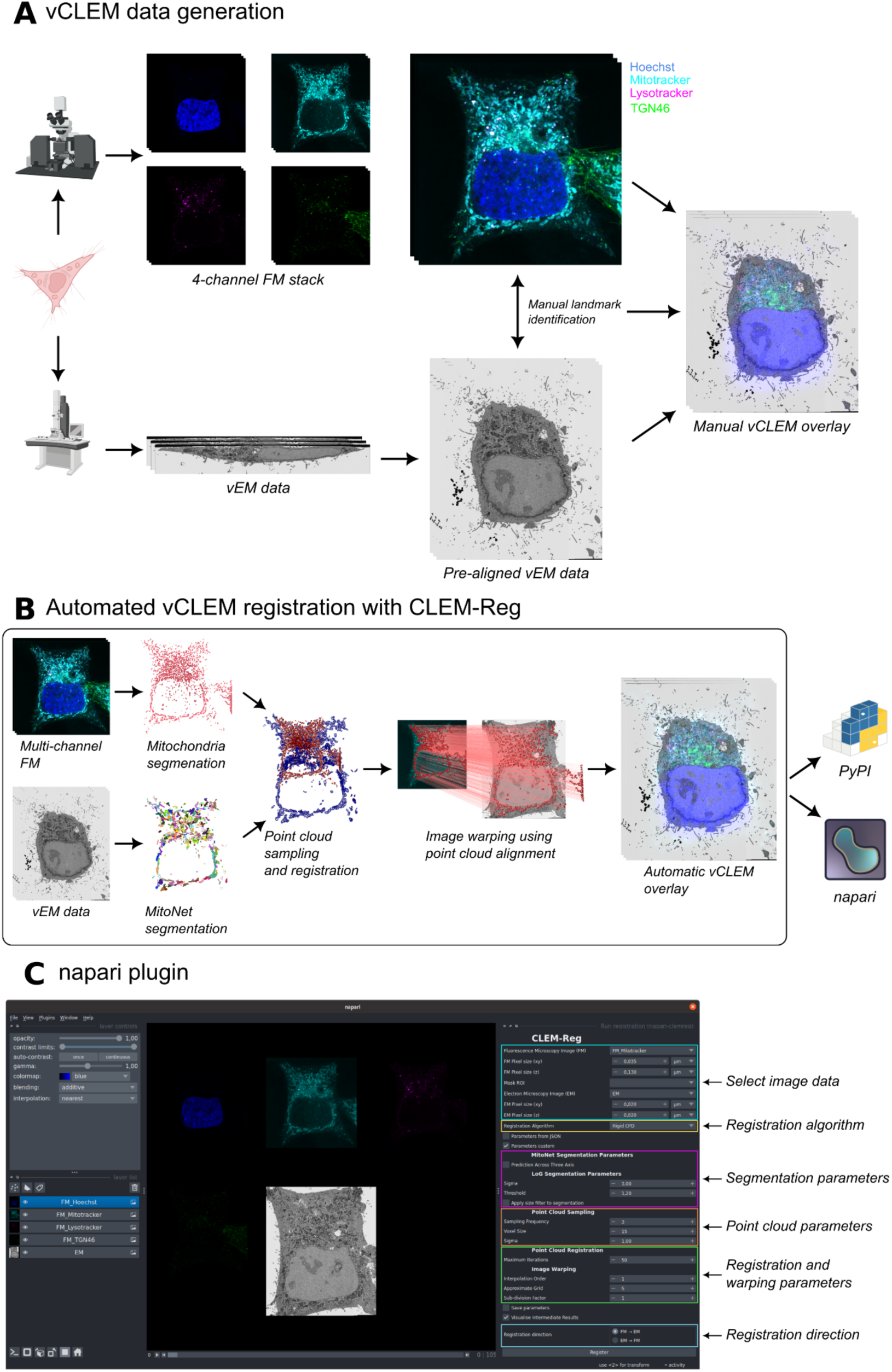
Volume CLEM data generation and CLEM-Reg algorithm. (A) Obtaining a vCLEM dataset consists of acquiring a FM and vEM image of the same sample. The two image stacks are manually aligned by identifying landmarks in both modalities and computing a transform to warp the FM stack onto the vEM data. (B) CLEM-Reg fully automates the registration step for vCLEM datasets by first segmenting mitochondria in both image modalities, sampling point clouds from these segmentations and registering them. Once registered, the point cloud alignment is used to warp the FM stack onto the vEM data. All data visualisations generated with napari. (C) The napari-clemreg plugin automatically registers vCLEM datasets with a single button click.

### The CLEM-Reg pipeline

CLEM-Reg automatically registers vCLEM data by segmenting mitochondria in FM and EM that are then used to generate point clouds, which are aligned with CPD (27,28), a state-of-the-art point cloud registration technique. The FM volume is then warped onto the EM volume using the found transformation (**Fig. 1B**). To aid adoption, CLEM-Reg is deployed as a plugin (“napari-clemreg”) for the napari image viewer (36), giving users the option for a single-click end-to-end operation, or to fine-tune or even entirely replace individual workflow steps, for example, importing segmentations from alternative sources (**Fig. 1C**).

### Segmenting internal landmarks

A promising approach to fully automate the CLEM alignment process is to automatically identify internal landmarks, speeding up the process and minimising opportunities for inadvertent subjective bias to occur. CLEM-Reg relies on the identification and segmentation of these common internal landmarks in both imaging modalities. For the purposes of this study, mitochondria were used as landmarks.

To obtain segmentations in FM, an algorithm based on combining a 3D Laplacian of Gaussian (LoG) filter with dynamic thresholding to account for signal degradation at deeper imaging levels was developed. The algorithm requires two parameters to be adjusted: kernel size of the LoG and the relative segmentation threshold. After obtaining an initial segmentation mask, spurious segmentations are removed with a size-based filter (**Fig. 2A**).

**Figure 2.**
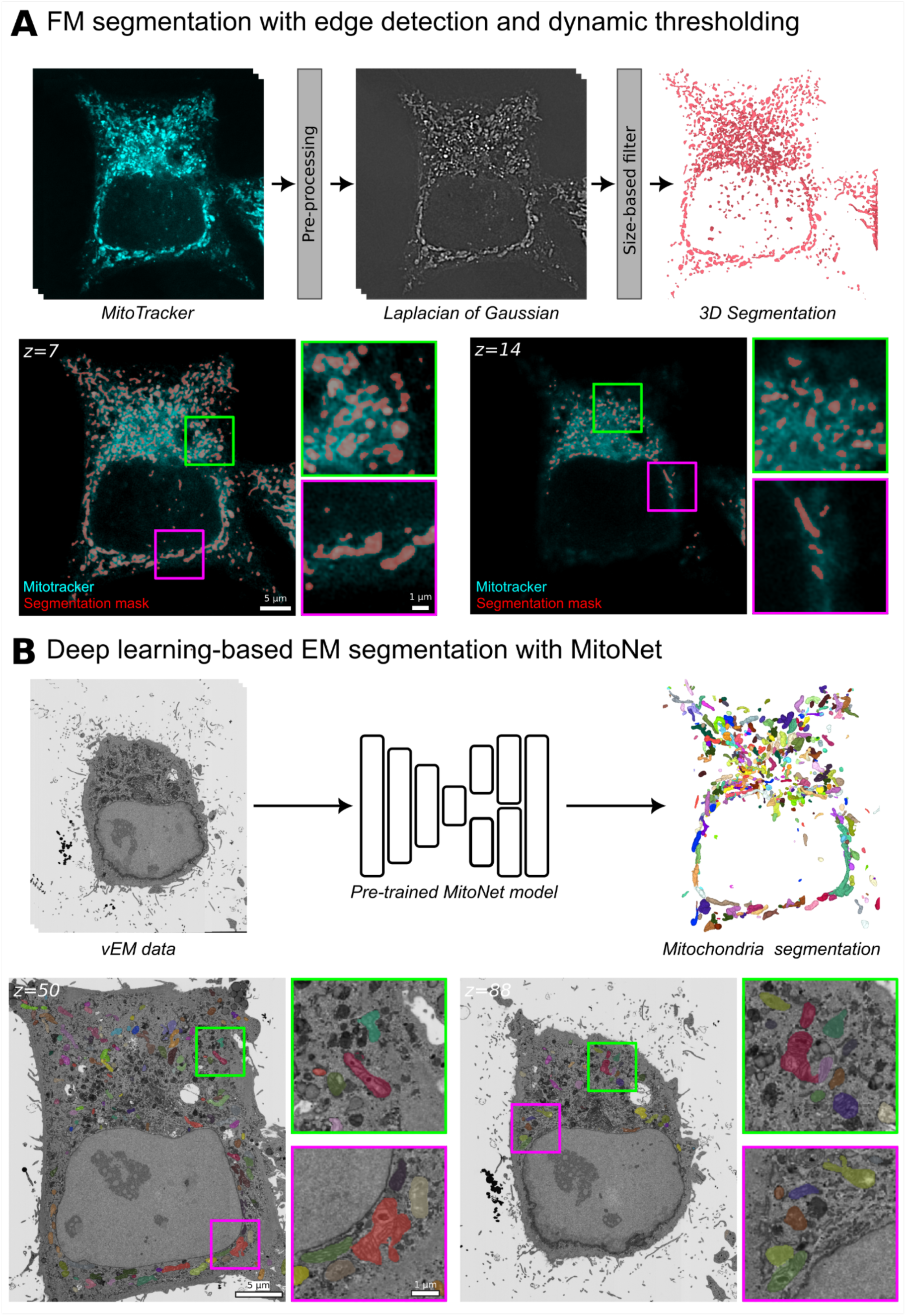
Mitochondria segmentation in FM and EM. (A) Mitochondria in the Mitotracker channel are segmented by applying a 3D Laplacian of Gaussian (LoG) filter to extract edges and dynamically thresholded to account for decreasing pixel intensity values as the imaging depth increases. To remove spurious mitochondria segmentations, a size-based filter is used. (B) Mitochondria in the vEM data are segmented with a pre-trained MitoNet (25) model. All visualisations generated with napari.

Mitochondria segmentations in EM are obtained with a pre-trained MitoNet (25) deep learning model which was found to perform well on FIB-SEM data out-of-the-box (**Fig. 2B**) and required slight preprocessing for SBF-SEM data (**Materials and Methods**). CLEM-Reg is however not restricted to using mitochondria segmentations and can be used with previously obtained EM segmentation masks of other structures or organelles (e.g. nuclear envelope).

### Generating modality-agnostic point clouds from internal landmarks and registration

The alignment between the FM and EM segmentations can be inferred by sampling 3D point clouds from the previously obtained segmentation masks. This reduces the computational load for large datasets and allows for mistakes in the segmentation to be ignored by using a probabilistic registration algorithm, such as CPD. The extraction of the coordinates of pixels from the exterior of the segmentation masks results in a 3D point cloud that captures its global architecture with an efficient representation due to the inherent sparsity of point clouds. The number of points in both point clouds depends on two parameters: binning and downsampling factor (**Fig. 3A**). Increasing any of these two parameters speeds up the registration, potentially by orders of magnitude. For instance, reducing the point sampling frequency from 1/16 to 1/256 with a fixed voxel size of 10 leads to a 19-fold decrease (from 33.7 min to 1.8 min) in registration time with no change in registration performance on EMPIAR-10819 (**Supp. Fig. 1)**.

**Figure 3.**
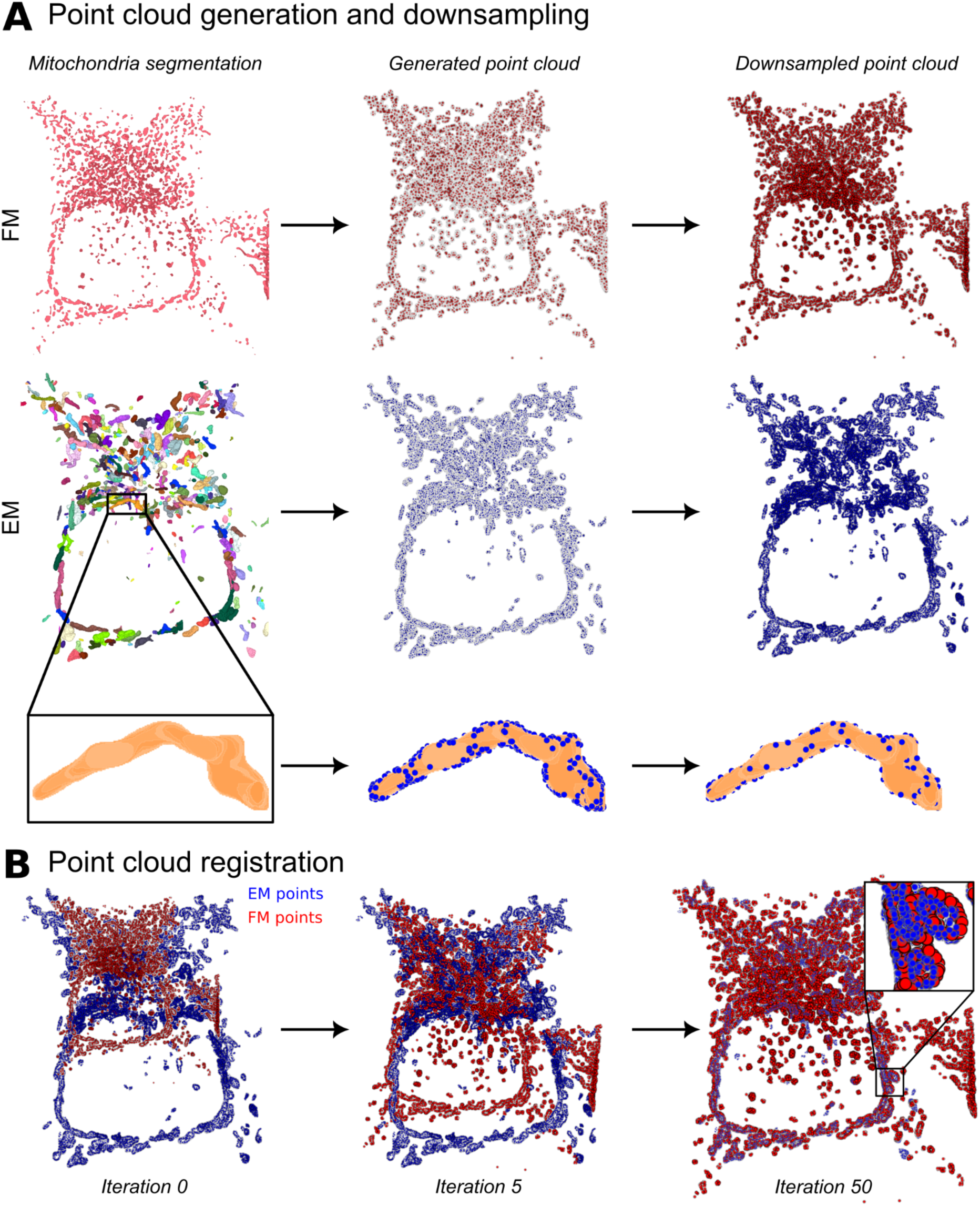
Point cloud generation and registration. (A) Point clouds are sampled on the surface of the 3D mitochondria segmentations in both FM and EM. To reduce the computational load and speed up the alignment time (**Supp. Fig. 1**), both point clouds are downsampled. (B) The point clouds are registered using rigid CPD (28) until convergence (50 iterations). All visualisations generated with napari.

After sampling, the point clouds are registered using either rigid CPD or non-linear Bayesian Coherent Point Drift (BCPD) (**Fig. 3B**) (27). Note that these probabilistic methods are necessary, as opposed to algorithms like Iterative Closest Point (ICP) (29), since the point sampling across modalities means there are no explicitly paired points. The choice of registration algorithm depends on the expected deformations between the FM and EM volumes, as well as computational constraints. In general, rigid CPD is faster and computationally less expensive than non-linear BCPD.

### Warping FM volume to obtain CLEM overlay

Once point clouds are registered, the found transformation is used to warp the FM volume onto the EM volume. This step is fast for rigid transformations but orders of magnitude slower for non-linear warping. CLEM-Reg implements 3D non-linear thin-plate spline warping (37) that uses the initially sampled and registered FM point clouds as control points. The runtime of the thin-plate spline warping depends on the interpolation order and size of the approximate grid. Since thin-plate spline warping is an expensive algorithm, CLEM-Reg also implements the option to sequentially warp sub-volumes. While this extends the runtime of the warping step, it reduces the required random access memory (RAM). Interestingly, overlays obtained with rigid alignment (**Fig. 4A-C**) outperformed non-linear alignment on the three benchmark datasets, assessed by metrics designed for the task.

**Figure 4.**
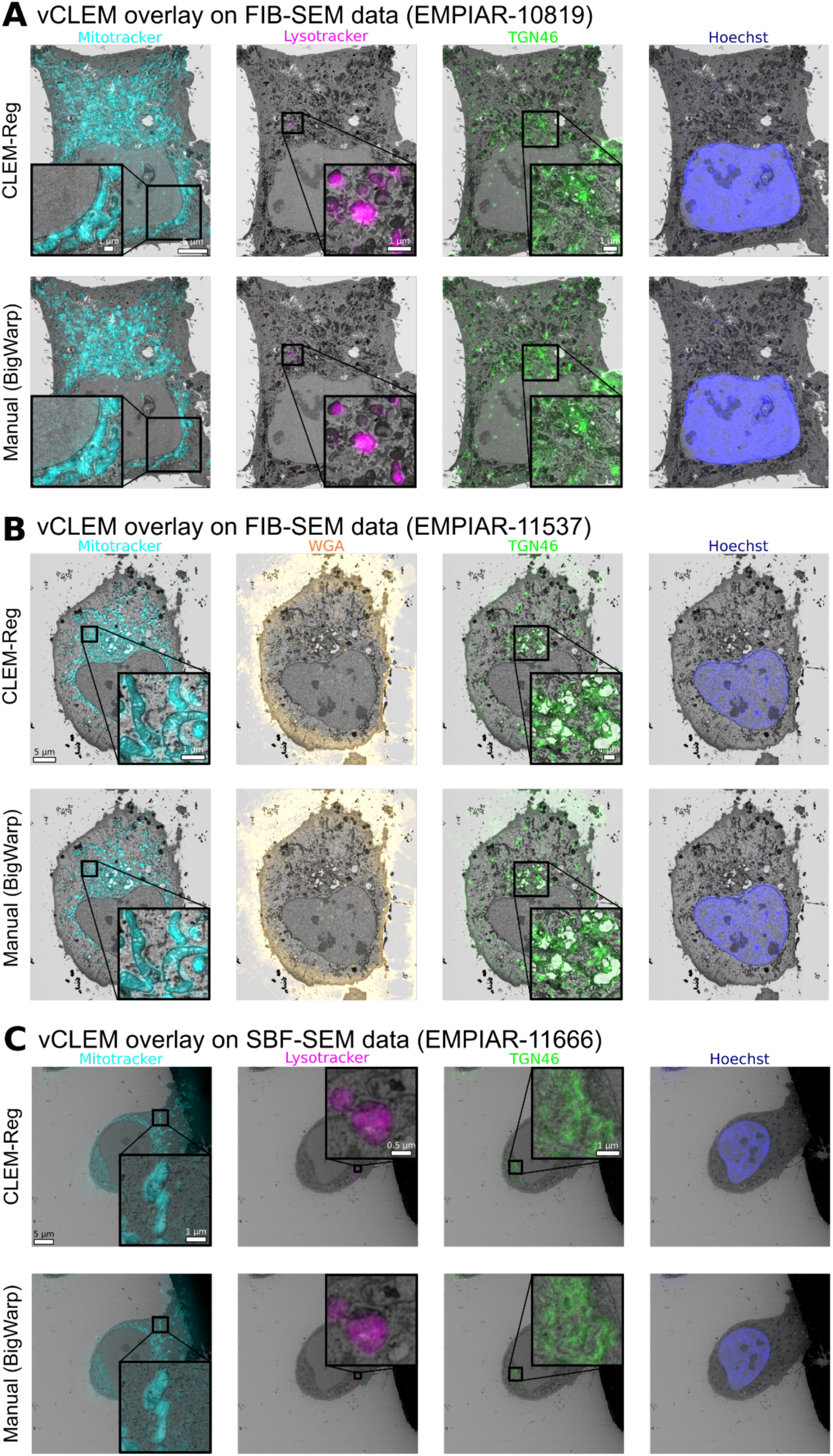
Comparing overlays obtained manually and with CLEM-Reg. CLEM-Reg overlays were obtained with rigid registration using the napari-clemreg plugin, while the manual overlays were obtained with affine registration using the BigWarp plugin in Fiji. (A) vCLEM overlays for EMPIAR-10819 dataset showing mitochondria (Mitotracker), lysosomes (Lysotracker), golgi apparatus (TGN46) and nucleus (Hoechst) with EM data acquired on FIB-SEM microscope. (B) vCLEM overlays for EMPIAR-11537 dataset showing Mitotracker, WGA, GFP-TGN46 and Hoechst with EM data acquired on FIB-SEM microscope. (C) vCLEM overlays for EMPIAR-11666 dataset showing Mitotracker, Lysotracker, GFP-TGN46 and Hoechst staining with EM data acquired on SBF-SEM microscope. Overlays generated with napari.

### Assessing CLEM-Reg performance against experts

Due to the lack of existing registration metrics for vCLEM alignment, two metrics were introduced to holistically assess registration performance: fluorescent signal overlap to EM structures and centroid distance between fluorescent signal and target structures.

For the correlation of fluorescence to FIB-SEM data, five target structures (lysosomes) were manually segmented in EM. The selected lysosomes varied in size (0.029 µm³ to 0.522 µm³) and were distributed across different areas of the cell, occupying both the centre and periphery (**Supp. Fig. 2A-B**).

To obtain CLEM-Reg overlays of the Lysotracker channel, off-target landmarks (mitochondria) were used to derive the warping matrix. This reduces bias that results from aligning fluorescence directly to presumed target structures. Registration with CLEM-Reg took 5.52 min (from starting registration to obtaining warped overlays for all channels) on a portable machine (**Materials and Methods**). Manual registration was performed with BigWarp by an expert with access to all fluorescent channels requiring around 2 h (**Fig. 5A**).

**Figure 5.**
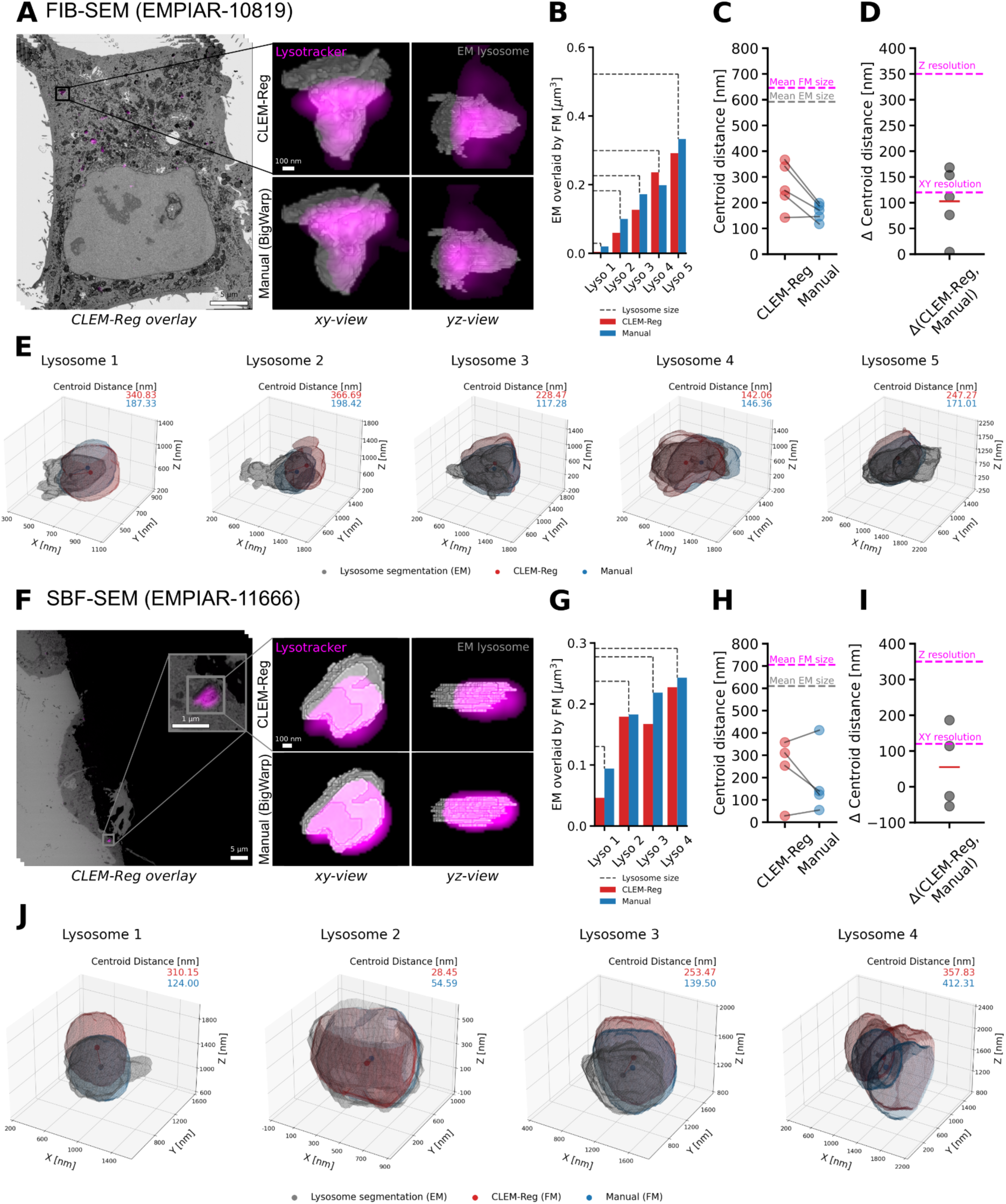
Comparing alignment results between CLEM-Reg and experts. (A, F) Lysotracker channels were overlaid to FIB-SEM (EMPIAR-10819) and SBF-SEM (EMPIAR-11666) data using mitochondria as off-target landmarks with CLEM-Reg. To quantify registration performance, five lysosomes were manually segmented throughout each EM volume. Corresponding segmentations in FM were obtained by segmenting the Lysotracker channel with Otsu’s method (38). (B, G) Volume of lysosomes in EM overlaid by FM signal was computed by intersecting EM and FM segmentations. (C, H) Centroid distances between EM segmentations and segmented Lysotracker signal in FM were computed with Euclidean distance. Mean size of FM and EM segmentations are shown in magenta and grey respectively. (D, I) The difference between Lysotracker signal overlaid manually and with CLEM-Reg was computed from previously found centroid distances. Mean difference in centroid distances is shown with a red horizontal line. The theoretical XY and Z resolution of the fluorescence microscope used is shown in magenta. (E, J) 3D visualisations of lysosome overlays were generated by obtaining meshes from segmentations of EM shown in grey, Lysotracker signal registered using BigWarp (Manual) shown in blue and CLEM-Reg shown in red. Visualisations generated with napari and matplotlib.

The volume of segmented lysosomes in EM overlaid by Lysotracker signal was computed by first segmenting the Lysotracker channel with Otsu thresholding (38) and then computing the intersected volume between both segmentations. It was found that all lysosomes segmented in EM were overlaid with the Lysotracker signal regardless of size (**Fig. 5B**). Notably, even the smallest lysosome with a volume of 0.029 µm³ was labelled with fluorescence, showing correct correlation well below the diffraction limit.

Next, centroid distances between segmented Lysotracker signal and lysosome segmentations were computed. Lysosomes were on average 2.62 times larger than average centroid distances obtained with CLEM-Reg, indicating unambiguous labelling (**Fig. 5C**). Average centroid distances obtained with CLEM-Reg were within 100.98 nm of centroid distances obtained from manual registration and thus below the theoretical resolution of the FM which, for the imaging system used, was 120 nm in XY and 350 nm in Z (**Fig. 5D)**. 3D visualisation overlays of segmentations and centroids of each lysosome are shown in **Fig. 5E**.

To assess the generalisability of CLEM-Reg to other EM modalities, performance was additionally assessed on a dataset acquired in SBF-SEM. CLEM-Reg overlays were obtained by registering mitochondria on a portable machine (**Materials and Methods**) requiring 2.44 min including image warping, while manual registration with BigWarp took around 2 h (**Fig. 5F**). A total of four lysosomes distributed across the cell were manually segmented for performance quantification (**Supp. Fig. 2C-F**).

CLEM-Reg overlaid Lysotracker signal to all four lysosomes in the SBF-SEM volume regardless of size (**Fig. 5G**). Lysosomes were on average 6.95 times larger than centroid distances obtained with CLEM-Reg, indicating unambiguous correlation between fluorescent signal and lysosomes (**Fig 5H**). On average, centroid distances in the overlay obtained with CLEM-Reg were within 54.88 nm of centroid distances obtained from manual registration. Notably, CLEM-Reg achieved smaller centroid distances compared to the manual registration on two lysosomes (**Fig 5I**). 3D visualisation overlays of segmentations and centroids of each lysosome are shown in **Fig. 5J**.

Overall, these results indicate that CLEM-Reg successfully automates vCLEM registration of both FIB-SEM and SBF-SEM data, correlating sub-micron structures with near expert-level accuracy while considerably reducing registration time. Crucially, registration with CLEM-Reg was performed using an off-target channel (Mitotracker) and assessed on an unseen target channel (Lysotracker) reducing potential biases arising from directly aligning target structures.

### CLEM-Reg plugin for napari

Napari is an open-source multi-dimensional image viewer for Python that allows third-parties to develop plugins which enable the addition of custom features and functionality tailored to their specific needs (36). The CLEM-Reg plugin was developed in napari due to the increasing adoption of Python in the bioimage analysis community and its seamless integration of state-of-the-art image processing packages and deep learning libraries. Furthermore, the “Points” and “Labels” layers support the visualisation of the point clouds and segmentations used in the CLEM-Reg workflow.

The plugin allows users to automatically register vCLEM datasets with a single click of a button. However, recognising that the ability for end-users to fine-tune their results can be crucial, the plugin also includes the option to execute and display intermediate steps of the full pipeline. Several different parameter configurations for a given step can thus be explored without needing to run the entire pipeline each time. (**Supp. Fig. 3**)

A range of features are included in the plugin, such as the option to delineate a corresponding ROI using the “Shapes” layer in napari, as well as the option to choose between rigid (CPD) and non-linear registration (BCPD). Parameters such as the FM segmentation settings, point cloud sampling density and the maximum number of iterations for the point cloud registration can be tuned. The chosen parameters can be saved and later re-used to ensure reproducibility. Overlays can be directly exported from napari, as well as the initial and transformed point clouds, for quality control purposes. Intermediate outputs like segmentation masks and sampled point clouds can be directly visualised in the napari viewer to aid troubleshooting. By utilising the “Layers” functionality of napari, it is also possible to inject intermediate results from external sources, for example, an FM segmentation method from a different plugin, or a pre-existing EM segmentation mask. The resulting “Labels” layer can then be set as an input to subsequent steps in CLEM-Reg. The results, usability and user-friendliness of the plugin were assessed on three vCLEM datasets (EMPIAR-10819, EMPIAR-11537 and EMPIAR-11666).

### Identifying TGN46-positive transport carriers with CLEM-Reg

To validate CLEM-Reg, a rare dynamic cellular process involving sub-micron organelles and transport carriers was studied. Despite being primarily localised to the TGN, a small subset of TGN46 (1 to 5%) rapidly cycles between TGN and plasma membrane. They exit the TGN in “CARriers of the TGN to the cell Surface” (CARTS), which play a role in transport of plasma membrane proteins (e.g. desmoglein-I, a key component of desmosomes) and secretion (e.g. lysozyme C, pancreatic adenocarcinoma upregulated factor (PAUF)) and recycle back to the TGN via endosomes (32–35). Leveraging the complementary information derived from vCLEM, unambiguous identification of the subset of endosomes and transport carriers engaged in the process of trafficking TGN46 between TGN and plasma membrane is demonstrated on two datasets (EMPIAR-10819 and EMPIAR-11537) with CLEM-Reg (**Fig. 6A-B**). Labelling accuracy of endosomes with GFP-TGN46 signal was quantitatively assessed on EMPIAR-11666 (**Supp. Fig. 4).**

**Figure 6.**
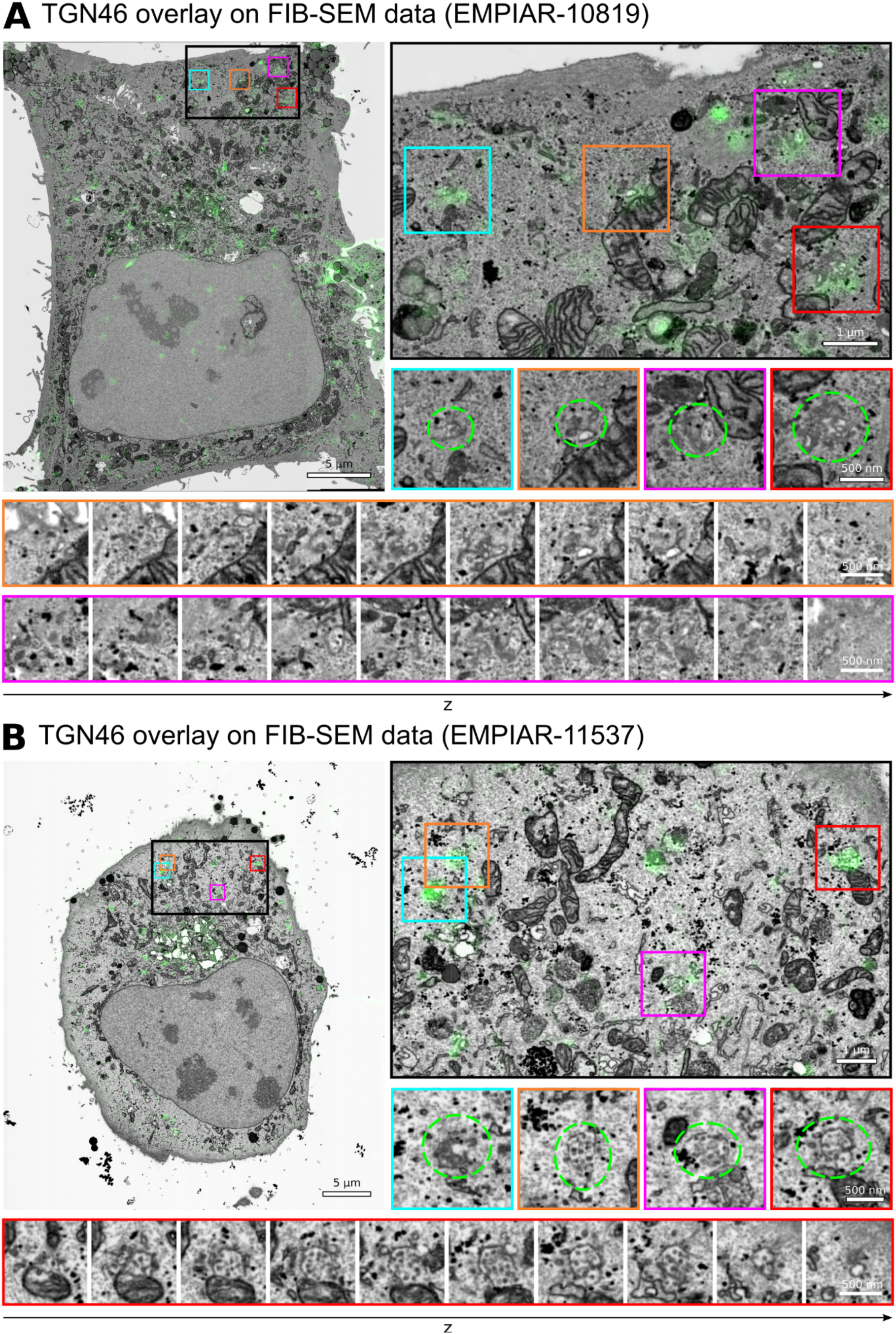
Identification of GFP-TGN46-positive endosomes and transport carriers. (A, B) In addition to the expected TGN localisation of GFP-TGN46, GFP-TGN46-positive endosomes and transport carriers were identified in EM guided by overlays obtained with CLEM-Reg. Structures of interest are circled in green and corresponding montages showing entire endosomes or transport carriers are displayed. Images in montages are shown with a spacing of 40 nm (orange) or 50 nm (magenta and red) in z. Full montages with 10 nm spacing in z can be found in **Supp. Fig. 5 and 6**.

By utilising off-target landmarks to derive the warping matrix (in this case, mitochondria), precise and accurate overlays of fluorescence to small target organelles is achieved whilst minimising subjective errors that result from aligning fluorescence directly to presumed target structures. Here, the overlays facilitated the identification of different morphological populations of TGN46-positive CARTS in different cellular locations: small vesicular structures tend to localise in the periphery, whereas complex membrane and vesicular structures are closer to the perinuclear region (**Suppl. Fig. 5 and 6**). Such functional labelling of organelles with sub-micron precision has the potential to provide vital mechanistic information in a range of biological processes.

## Discussion

This study introduces CLEM-Reg, an automated and unbiased end-to-end vCLEM registration framework. It can be easily accessed by end-users through a dedicated plugin for napari. The proposed registration approach relies on extracting landmarks with a pattern recognition approach that includes deep learning. After segmenting internal landmarks in FM and EM, point clouds are sampled and preprocessed to obtain a memory-efficient, modality-agnostic representation. By using state-of-the-art point cloud registration methods, a mapping between the two image volumes is found, with which the FM acquisition is warped onto the EM volume to obtain a final CLEM overlay. CLEM-Reg drastically reduces registration time to a few minutes and achieves near expert-level performance on three benchmark CLEM datasets.

CLEM-Reg’s potential in driving new biological insights is also demonstrated. Correlation of small punctate structures, like the transport carriers and endosomes studied herein, is challenging due to the lack of characteristic morphologies. Thus, accurate correlation requires precise and unbiased landmark-based alignment of off-target structures which CLEM-Reg automatically delivers across the whole cell volume. Wakana et al studied TGN46 carriers in 2D using permeabilised cells (35). CLEM-Reg however enables identification of TGN46 carriers in 3D across whole cells with intact membranes. Future work could build on these results by studying morphological differences of transport carriers in their native state between healthy and pathological cells. Since all registered datasets have been publicly deposited, others interested in TGN46 trafficking can mine these data to more completely study instances of TGN46-positive CARTS. Overall, these results highlight a broader concept for underpinning structure-function studies with vCLEM.

Nevertheless, certain limitations remain in regard to the automated organelle segmentation in vEM which directly impacts the alignment performance (**Supp. Fig. 7**). While MitoNet was shown to perform well on the three vCLEM datasets shown in this study, this cannot be guaranteed for all vEM datasets. For instance, to obtain mitochondria segmentations throughout the SBF-SEM dataset, preprocessing with contrast-limited adaptive histogram equalisation was required (**Materials and Methods**). This is likely due to the imbalanced dataset used to train MitoNet with 75.6% of data acquired on FIB-SEM and only 5.2% on SBF-SEM microscopes (25). One approach to address this potential limitation is to make use of transfer learning, a deep learning method in which already trained models are briefly retrained on a smaller dataset to improve performance. While training deep neural networks previously required access to GPUs and fluency in programming languages such as Python, open-source projects like ZeroCostDL4Mic (39), DeepImageJ (40), Imjoy (41) or DL4MicEverywhere (42) are rapidly removing these barriers by offering frameworks that leverage freely available computing resources for training, enabling GUI-driven interaction with a range of deep learning architectures. Open source and community-driven model libraries such as the Bioimage Model Zoo (43), which allow easy sharing of pre-trained deep learning models, are another important resource in this context.

CLEM-Reg includes the option to perform non-linear alignment of volumes using BCPD and 3D thin-plate-spline warping. However, rigid registration applied to a FIB-SEM benchmark dataset outperforms non-linear registration and also runs orders of magnitude faster (2.17 min. vs. 205 min) on a portable machine (**Materials and Methods**). A possible explanation for this counterintuitive finding is that better performance can be achieved by restricting the degrees of freedom, since the point-cloud data from the two modalities are inherently noisy and do not perfectly match. Thus, as noise increases due to factors like segmentation errors, non-isotropic pixel-sizes in the FM volume or point sampling, the non-linear registration approach is more prone to converging to non-optimal solutions due to the larger parameter space. Rigid registration restricts the degrees of freedom and thus the parameter space which facilitates convergence towards an optimal solution.

The size of bioimage datasets can also cause challenges for automated alignment. More specifically, downsampling of the FM and EM datasets is generally required prior to execution. Despite this, downsampling may still need to be improved for very large datasets. To circumvent this limitation, the use of next-generation file formats (44) and tools such as Dask (45) could enable the computation of datasets that exceed available memory by chunking, thus allowing users to run the CLEM-Reg workflow using full resolution data on conventional computers without encountering memory issues. Additionally, the CLEM-Reg workflow relies heavily on two packages, Scipy (46) and scikit-image (47), for tasks such as FM segmentation and point sampling of the FM and EM segmentation. However, these packages do not natively provide GPU acceleration, thus do not benefit from available compute resources for users possessing GPUs. As such, future work to incorporate GPU acceleration into the CLEM-Reg workflow using Python libraries such as Cupy (48) or clEsperanto (49) could reduce processing time. This optimisation can significantly accelerate multiple steps within the CLEM-Reg workflow.

While CLEM-Reg’s ability to register vEM techniques that image the block face is demonstrated, other vEM methods like array tomography and serial section TEM could be used given a sufficiently accurate section-to-section alignment. However, robust alignment of consecutive EM sections is challenging, due to the non-linear deformations associated with manual sectioning and imaging, in particular by TEM. Currently, the alignment is mostly done manually, with each section finely aligned to the previous section using manually placed landmarks on adjacent sections or manually applying linear or non-linear transforms to each image. This process can introduce bias and is extremely lengthy, requiring days of an expert’s time, and therefore beyond the scope of this paper. We speculate that methods based on registering segmented landmarks, such as CLEM-Reg, could form part of an iterative pipeline to improve this section-to-section alignment by registering objects in consecutive slices.

While CLEM-Reg has been demonstrated to register single-cell scale data via the segmentation of mitochondria, the core of the alignment workflow is agnostic to the landmarks being segmented. As such, the method could be used at different scales or across different modalities as long as certain conditions are met. The main conditions are the existence of a segmentation algorithm for the same structure in both modalities, and that those structures are numerous and distributed throughout the volume to be aligned. For example, nucleus segmentation algorithms in EM and FM could be used to derive point clouds capable of registering tissue-scale volumes for vCLEM, or nucleus segmentation in X-ray microscopy could be aligned to either EM or FM (or both). With the rapid development of novel AI-based segmentation methods (50), it is expected that the range of suitable target structures will continue to grow. These developments hold the promise of broadening the applicability of CLEM-Reg to other multimodal registration tasks beyond vCLEM.

## Materials and Methods

### Cell model

Human cervical cancer epithelial cells (HeLa) were obtained from the Cell Services Science Technology Platform at The Francis Crick Institute and originated from the American Type Culture Collection (ATCC; CCL-2).

### CLEM data acquisition

HeLa cells were maintained in Dulbecco’s Modified Eagle Medium (Gibco) supplemented with 10% foetal bovine serum at 37°C and 5% CO2. 150,000 cells were seeded in 35 mm photo-etched glass-bottom dishes (MaTtek Corp). At 24 h, cells were transfected with 0.1 µg of GFP-TGN46 construct per dish, using Lipofectamine LTX and PLUS reagent (Invitrogen) in Opti-MEM medium (Gibco) as recommended by the manufacturer. 300 µl of transfection mix were added to 2 ml antibiotic-free medium and incubated overnight. At 36 h, cells were stained with Lysotracker red (100 nM) or Wheat Germ Agglutinin (WGA) Alexa Fluor 594 (5 µg/ml) and Mitotracker deep red FM (200 nM) for 10 min in HBSS (WGA) or 30 min in DMEM (Lysotracker and Mitotracker). Hoechst 33342 (1 µg/µl) was added in the last 5 min of the tracker incubation. All probes were from Molecular Probes (Thermo Fisher Scientific). Cells were then washed 3 times with 0.1 M phosphate buffer (PB) pH 7.4 and fixed with 4% (v/v) formaldehyde (Taab Laboratory Equipment Ltd) in 0.1 M PB pH 7.4 for 15 min. Cells were washed twice and imaged in 0.1 M PB on an AxioObserver 7 LSM900 with Airyscan 2 microscope with Zen 3.1 software (Carl Zeiss Ltd). Cells were first mapped with a 10x objective (NA 0.3) using brightfield microscopy to determine their position on the grid and tile scans were generated. The cells of interest were then imaged at high resolution in Airyscan mode with a 63x oil objective (NA 1.4). Smart setup was used to set up the imaging conditions. A sequential scan of each channel was used to limit crosstalk and z-stacks were acquired throughout the whole volume of the cells.

The samples were then processed using a Pelco BioWave Pro+ microwave (Ted Pella Inc, Redding, USA) following a protocol adapted from the National Centre for Microscopy and Imaging Research protocol (51). Each step was performed in the Biowave, except for the phosphate buffer and water wash steps, which consisted of two washes on the bench followed by two washes in the Biowave without vacuum (at 250 W for 40 s). All the chemical incubations were performed in the Biowave for 14 min under vacuum in 2 min cycles alternating with/without 100 W power. The SteadyTemp plate was set to 21°C unless otherwise stated. In brief, the samples were fixed again in 2.5% (v/v) glutaraldehyde (TAAB) / 4% (v/v) formaldehyde in 0.1 M PB. The cells were then stained with 2% (v/v) osmium tetroxide (TAAB) / 1.5% (v/v) potassium ferricyanide (Sigma), incubated in 1% (w/v) thiocarbohydrazide (Sigma) with SteadyTemp plate set to 40°C, and further stained with 2% osmium tetroxide in ddH2O (w/v). The cells were then incubated in 1% aqueous uranyl acetate (Agar Scientific) with a SteadyTemp plate set to 40°C, and then washed in dH2O with SteadyTemp set to 40°C. Samples were then stained with Walton’s lead aspartate with SteadyTemp set to 50°C, and dehydrated in a graded ethanol series (70%, 90%, and 100%, twice each), at 250 W for 40 s without vacuum. Exchange into Durcupan ACM® resin (Sigma) was performed in 50% resin in ethanol, followed by 4 pure Durcupan steps, at 250 W for 3 min, with vacuum cycling (on/off at 30 s intervals), before polymerisation at 60°C for 48 h.

Focused ion beam scanning electron microscopy (FIB-SEM) data was collected using a Crossbeam 540 FIB-SEM with Atlas 5 for 3D tomography acquisition (Carl Zeiss Ltd). A segment of the resin-embedded cell monolayer containing the cell of interest was trimmed out, and the coverslip was removed using liquid nitrogen prior to mounting on a standard 12.7 mm SEM stub using silver paint and coating with a 10 nm layer of platinum. The region of interest was relocated by briefly imaging through the platinum coating at an accelerating voltage of 10 kV and correlating to previously acquired fluorescence microscopy images. On completion of preparation of the target ROI for Atlas-based milling and tracking, images were acquired at 5 nm isotropic resolution throughout the cell of interest, using a 6 µs dwell time. During acquisition, the SEM was operated at an accelerating voltage of 1.5 kV with 1.5 nA current. The EsB detector was used with a grid voltage of 1.2 kV. Ion beam milling was performed at an accelerating voltage of 30 kV and a current of 700 pA.

For serial block face scanning electron microscopy (SBF-SEM), the block was trimmed to a small trapezoid, excised from the resin block, and attached to an SBF-SEM specimen holder using conductive epoxy resin. Prior to the commencement of an SBF-SEM imaging run, the sample was coated with a 2 nm layer of platinum to further enhance conductivity. SBF-SEM data was collected using a 3View2XP (Gatan) attached to a Sigma VP SEM (Carl Zeiss Ltd). Inverted backscattered electron images were acquired through the entire extent of the region of interest. For each of the 133 consecutive 50 nm slices needed to image the cell in its whole volume, a low resolution overview image (horizontal frame width 150.58 µm; pixel size of 75 nm; 2 µs dwell time) and a high resolution image of the cell of interest (horizontal frame width 63.13 µm; pixel size of 7 nm; using a 3 µs dwell time) were acquired. The SEM was operated in high vacuum with focal charge compensation on. The 30 µm aperture was used, at an accelerating voltage of 2 kV.

### Pre-processing for CLEM registration

The Airyscan data was first processed in Zen software using the z-stack alignment tool to correct z-shift. The settings used were the highest quality, translation and linear interpolation, with the Mitotracker channel as a reference. The .czi file was then opened in Fiji (52) and saved as .tif. For EMPIAR-11666, the Mitotracker channel was further processed using Yen’s thresholding method (53) to remove overexposed punctate artefacts affecting FM segmentation.

After initial registration with template matching by normalised cross-correlation in Fiji (https://sites.google.com/site/qingzongtseng/template-matching-ij-plugin), the FIB-SEM images (EMPIAR-10819 and EMPIAR-11537) were contrast normalised as required across the entire stack and converted to 8-bit grayscale. To fine-tune image registration, the alignment to the median smoothed template method was applied (54). The aligned XY 5 nm FIB-SEM stack was opened in Fiji and resliced from top to bottom (i.e., to XZ orientation), and rotated to roughly match the previously acquired Airyscan data. Finally, the EM volume was binned by a factor of 2 and 4 in X, Y and Z using Fiji resulting in an isotropic voxel-size of 10 and 20 nm respectively.

For the SBF-SEM data (EMPIAR-11666), only minor adjustments in image alignment were needed and were done using TrakEM2 in Fiji (55). Finally, the EM volume was binned by a factor of 2 and 4 in XY to obtain a voxel-size of 14×14×50 nm and 28*×*28*×*50 nm respectively.

### Computational resources

The algorithms described here were primarily developed and tested on a portable machine running on a Linux distribution with the following specifications: 64 GB RAM, Intel Core i7-11850H 2.50GHz*×*16 CPU and GeForce RTX 3080 GPU. An additional workstation with the following specs was also used for testing: 256 GB RAM, Dual Intel Xeon (14 CPU each) 2.6GHz, Quadro M600 16GB GPU.

### FM segmentation with thresholded Laplacian of Gaussian

To ensure the FM pixel values were in the range of 0 and 1, min-max normalisation of the following formula was used, where *x* is the FM volume, *x_min_* is the lowest pixel value in the volume and *x_max_* is the largest pixel value in the volume and *x_scaled_* is the resulting scaled volume 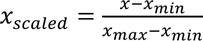. In the subsequent step, the scaled FM volume undergoes a LoG filtering process, requiring one tunable parameter σ. This process involves convolving a 1D Difference of Gaussians (DoG) across the X, Y and Z planes of the volume. Marr and Hildreth (56) demonstrated that a reasonable approximation of LoG can be attained by maintaining a ratio of 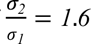 when applying the DoG. A value of σ*_1_* σ = *3*.*0* was used for EMPIAR-10819 and EMPIAR-11537, thus after applying the LoG ratio, σ*_1_* = *3*.*0* and σ*_2_* = *4*.*8*. A dynamic threshold based on a relative user-defined threshold and the mean intensity of a given slice was then applied to each image in the resulting volume to account for inconsistencies arising due to attenuation of the signal at deeper imaging levels. The relative threshold was set to *T* = *1*.*2* for EMPIAR-10819 and EMPIAR-11537. For EMPIAR-11666, the following parameters were used: σ = *2*.*2* and *T* = *1*.*3*. Lastly, to reduce the number of spurious segmentations, a size-based filter was then applied. This was achieved by using a 3D connected components algorithm from the Python package cc3d 3.2.1 (57) which removes components below a user-defined size threshold with a default value set to remove all segmentations with a size below the 5th and above the 95th percentile.

### EM segmentation with MitoNet

The mitochondria within the EM volume were segmented using a pre-trained MitoNet deep learning model (58). The MitoNet model using its default parameters was applied to EM volumes in EMPIAR-10819 and EMPIAR-11537 without preprocessing, resulting in instance segmentation of the mitochondria. To improve mitochondrial segmentation of EMPIAR-11666, the aligned stack was processed for contrast-limited adaptive histogram equalisation (59) using the Enhance Local Contrast (CLAHE) plugin in Fiji, using the following parameters: blocksize **=** 127; histogram bins **=** 256; maximum slope **=** 2, 3 and 10; mask **=** “*None*”; fast. Each CLAHE stack (from 3 maximum slope values) was segmented using MitoNet and the masks were merged by summation before point cloud sampling.

### Point cloud sampling of the EM and FM segmentation

The method employed for producing point clouds from the FM and EM volumes uses the same standardised procedure. Firstly, the FM and EM segmentation masks were resampled to obtain isotropic voxels (EMPIAR-10819 and EMPIAR-11537: 20 nm; EMPIAR-11666: 50 nm). Then, a Canny edge filter (60) with a default value of σ = 1.0 was applied to each binary segmentation slice. This process ensures that only points on the edge of the outer mitochondrial membrane are randomly sampled. Then, every pixel along the membrane was identified as a potential point to be used within the point cloud, but due to the computationally expensive nature of point cloud registration algorithms, subsequent downsampling was carried out.

### Point cloud downsampling and registration with CPD and BCPD

After sampling, the point clouds were downsampled, binned and filtered by removing statistical outliers to reduce memory requirements and noise. Downsampling of the point clouds was performed by a uniform downsampling function which takes the point cloud and user-defined sampling frequency *1*/*k* as the input and samples every *k*^-*ℎ*^ point in the point cloud, starting with the *0*^-*ℎ*^ point, then the *k*^-*ℎ*^ point, then the *k* + *k*^-*ℎ*^ point and so on. A value of *k* = *30* was used for all datasets. Following the initial downsampling, a binning step was used which employs a regular voxel grid to create a downsampled point cloud from the input point cloud. There are two primary steps of the binning starting with an initial partitioning of the point cloud into a set of voxels. Subsequently, each occupied voxel generates exactly one point through the computation of the average of all the points enclosed within it, resulting in a binned version of the original point cloud. A voxel-size of *s* = *15* was used for all datasets. For registration, the Python package probreg 0.3.6 (61) was used, as it implements both the CPD (28) and BCPD (27) algorithm with a unified API. For all datasets, rigid CPD with 50 iterations was used to obtain globally registered point clouds of the FM to the EM. The found transformation matrix is then used for the warping of the image volumes.

### Warping of image volumes

The transformation matrix found from the global point cloud registration was applied to the source image volume using the affine transform implementation provided by SciPy 1.10.1 (46), a powerful open-source Python library that provides a range of functions for mathematics, science, and engineering. This implementation makes use of inverse projection to interpolate the intensity values of the warped image thus requiring an inverted transformation matrix which is found with NumPy’s 1.24.2 (62) linear algebra module. Interpolation of the warped image was achieved with higher-order spline interpolation. Non-linear image warping was achieved with thin-plate spline deformation which finds a mapping between a set of control-points *f*: (*x*, *y*, *z*) → (*x′*, *y′*, *z′*) such that the deformed source points match the target points as closely as possible while ensuring that the image between these control-points remains smooth. As there are no open-source implementations for volumetric thin-plate spline warping available for Python, a custom script was implemented based on (37). To reduce computational overhead, the FM volume was nonlinearly warped as eight individual chunks. After warping, FM volumes are resampled to map to the EM space.

### Benchmarking of CLEM-Reg against experts

For expert manual registration, image stacks from FM and EM were manually aligned to each other using the BigWarp plugin of the Fiji framework, with the EM stack set as “target” and the FM stack as “moving” dataset. Corresponding points were placed on mitochondria on both datasets and throughout the whole volume of the cell. An affine transformation was applied to the FM data and the transformed dataset was merged with the EM data to produce the final overlay.

Due to the inherent differences in appearance between the two imaging modalities, direct intensity-based quantification of the registration performance is not possible. Therefore, two metrics were introduced to assess registration performance: fluorescent signal overlap to EM structures and centroid distance between fluorescent signal and target structures.

Firstly, target structures (lysosomes in EMPIAR-10819 and EMPIAR-11666 and endosomes in EMPIAR-11537) were manually segmented in 3D in the EM volume in TrakEM2 in Fiji. These lysosomes or endosomes were then cropped with a bounding box encompassing the fluorescent signal from the corresponding manual- or CLEM-Reg-warped FM volumes and then segmented using the Otsu thresholding method (38) to generate the correlating FM segmentation.

Segmentations were transformed from pixel space to real space by applying appropriate scaling. The intersected volume between fluorescent and EM segmentations was then computed. Centroid distances were obtained by first constructing meshes from segmentations using marching cubes. Then, centroids were determined on meshes derived from fluorescent and EM segmentations and their Euclidean distance calculated. The size of segmented target structures (lysosomes or endosomes) was estimated by computing characteristic length scales *L* from the volume of the segmented target structures *V* with 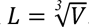.

### Plugin development and deployment

The plugin was developed for Python 3.9 and napari 0.4.17 (36). It integrates the CLEM-Reg workflow in the form of a single widget running end-to-end and as a set of individual widgets, each carrying out an individual step (split registration functionality). The EMPIAR-10819 dataset can directly be loaded as sample data from the plug-in. To distribute the plugin and make it accessible from the napari interface, the project is packaged using the cookiecutter plugin template provided by napari. The plugin is available for installation via the package installer for Python (pip) in the command line under the project name “napari-clemreg”. The plugin can also be installed via the built-in plugin menu which lists all plugins currently available on napari-hub. The source-code is available as a GitHub repository alongside a detailed project description, installation instructions and user guide.

## DATA AND CODE AVAILABILITY

The datasets used in this study have been deposited at EMPIAR and Biostudies. EMPIAR-10819: EM (https://www.ebi.ac.uk/empiar/EMPIAR-10819/) FM (https://www.ebi.ac.uk/biostudies/bioimages/studies/S-BSST707). EMPIAR-11537: EM (https://www.ebi.ac.uk/empiar/EMPIAR-11537/) FM (https://www.ebi.ac.uk/biostudies/bioimages/studies/S-BSST1075). EMPIAR-11666: EM (https://www.ebi.ac.uk/empiar/EMPIAR-11666/) FM (https://www.ebi.ac.uk/biostudies/bioimages/studies/S-BSST1175). Code for CLEM-Reg (under MIT licence) is available on GitHub (https://github.com/krentzd/napari-clemreg).

## ACKNOWLEDGEMENTS

This work was supported by The Francis Crick Institute which receives its core funding from Cancer Research UK (CC1076), the UK Medical Research Council (CC1076), and the Wellcome Trust (CC1076). D.K. was funded by The Francis Crick Institute, the Pasteur-Paris University International doctoral program (PPU), the INCEPTION program and a Fondation pour la Recherche Médicale (FRM) Fin de Thèse (FDT) grant. R.F.L. was supported by a Medical Research Council Skills development fellowship (MR/T027924/1). R.H. is supported by the Gulbenkian Foundation (Fundação Calouste Gulbenkian) and received funding from the European Research Council (ERC) under the European Union’s Horizon 2020 research and innovation program (grant agreement number 101001332), the European Union through the Horizon Europe program (AI4LIFE project with grant agreement 101057970-AI4LIFE, and RT-SuperES project with grant agreement 101099654-RT-SuperES to R.H.), the European Molecular Biology Organization (EMBO) Installation Grant (EMBO-2020-IG4734),the Chan Zuckerberg Initiative Visual Proteomics Grant (vpi-0000000044 with doi:10.37921/743590vtudfp) and a Chan Zuckerberg Initiative Essential Open Source Software for Science (EOSS6-0000000260). Views and opinions expressed are those of the authors only and do not necessarily reflect those of the European Union. Neither the European Union nor the granting authority can be held responsible for them. Matt Russell (Electron Microscopy Science Technology Platform, The Francis Crick Institute and Centre for Ultrastructural Imaging, King’s College London) collected preliminary CLEM data used in testing.

## CONTRIBUTIONS

DK, RFL, RH, LMC and MLJ developed the initial idea for this project. DK conceived CLEM-Reg and directed this project. DK, ME, MCD, MLJ and LMC wrote the manuscript. DK and ME developed the source-code. CS created a Docker file. RFL collected preliminary CLEM data. MCD and CJP collected the benchmark CLEM datasets (MCD: FM and SBF-SEM, CJP: FIB-SEM). MCD performed manual registrations, segmentations and user testing. DK, ME, MCD, RFL, RH, LMC and MLJ contributed to the revisions of the manuscript.

