## Supplementary Material for "CLEM-Reg: An automated point cloud based registration algorithm for correlative light and volume electron microscopy"

### Supplementary Notes

**Benchmarking point sampling frequency and binning.** Reducing the number of sampled points from the identified landmarks reduces the registration time. Two parameters control point sampling: sampling frequency and binning. The first of these parameters removes every k$^{th}$ point from the point cloud while the second parameter bins the point cloud by a given voxel size. The normalised root mean squared error (NRMSE) between CLEM-Reg and manual overlays was computed on Mitotracker and Lysotracker/GFP-TGN46 channels as a proxy for registration performance. We find that reducing the point sampling frequency from 1/16 to 1/256 with a fixed voxel size leads to a 19-fold decrease (from 33.7 min to 1.8 min) in registration time with no change in registration error. (**Supp. Fig. 1**). This shows that by tuning the number of points used to represent landmarks, drastic reductions in registration time can be obtained without impacting registration accuracy.

**Performance quantification on endsomes.** Firstly, target structures (endosomes in EMPIAR-11537) were manually segmented in 3D in the EM volume in TrakEM2 in Fiji. These endosomes were then cropped with a bounding box from the corresponding manual- or CLEM-Reg-warped FM volumes and then segmented using the Otsu thresholding method to generate the correlating FM segmentation. Segmentations were transformed from pixel space to real space by applying appropriate scaling. The intersected volume between fluorescent and EM segmentations was then computed. Centroid distances were obtained by first constructing meshes from segmentations using marching cubes. Then, centroids were determined on meshes derived from fluorescent and EM segmentations and their Euclidean distance calculated. The size of segmented endosomes was estimated by computing characteristic length scales $L$ from the endosome volume $V$ with $L = \sqrt[3]{V}$ (**Supp. Fig. 4**).

**Assessing the robustness of CLEM-Reg to segmentation errors.** The CLEM-Reg algorithm requires identification of landmark structures in both FM and EM to obtain an alignment between image volumes. While the segmentation algorithms used in the context of the benchmark data perform well, the robustness of the segmentation algorithms cannot be guaranteed. We reasoned that the deep learning-based segmentation approach using MitoNet would be more susceptible to performance degradation on new EM image volumes and therefore tested the robustness of CLEM-Reg to random loss of mitochondria segmentations across the whole EM volume. The normalised root mean squared error (NRMSE) between CLEM-Reg and manual overlays was computed on Mitotracker and Lysotracker/GFP-TGN-46 channels as a proxy for registration performance. We find that the registration performance is constant up to a loss of around 40% of mitochondria (**Supp. Fig. 7A**). We also assessed the impact of segmentation errors in different areas of the EM volume and found that the registration performance is more sensitive to the loss of peripheral landmarks, as opposed to losing landmarks in the central region of the EM volume (**Supp. Fig. 7B**).

### Supplementary Figures

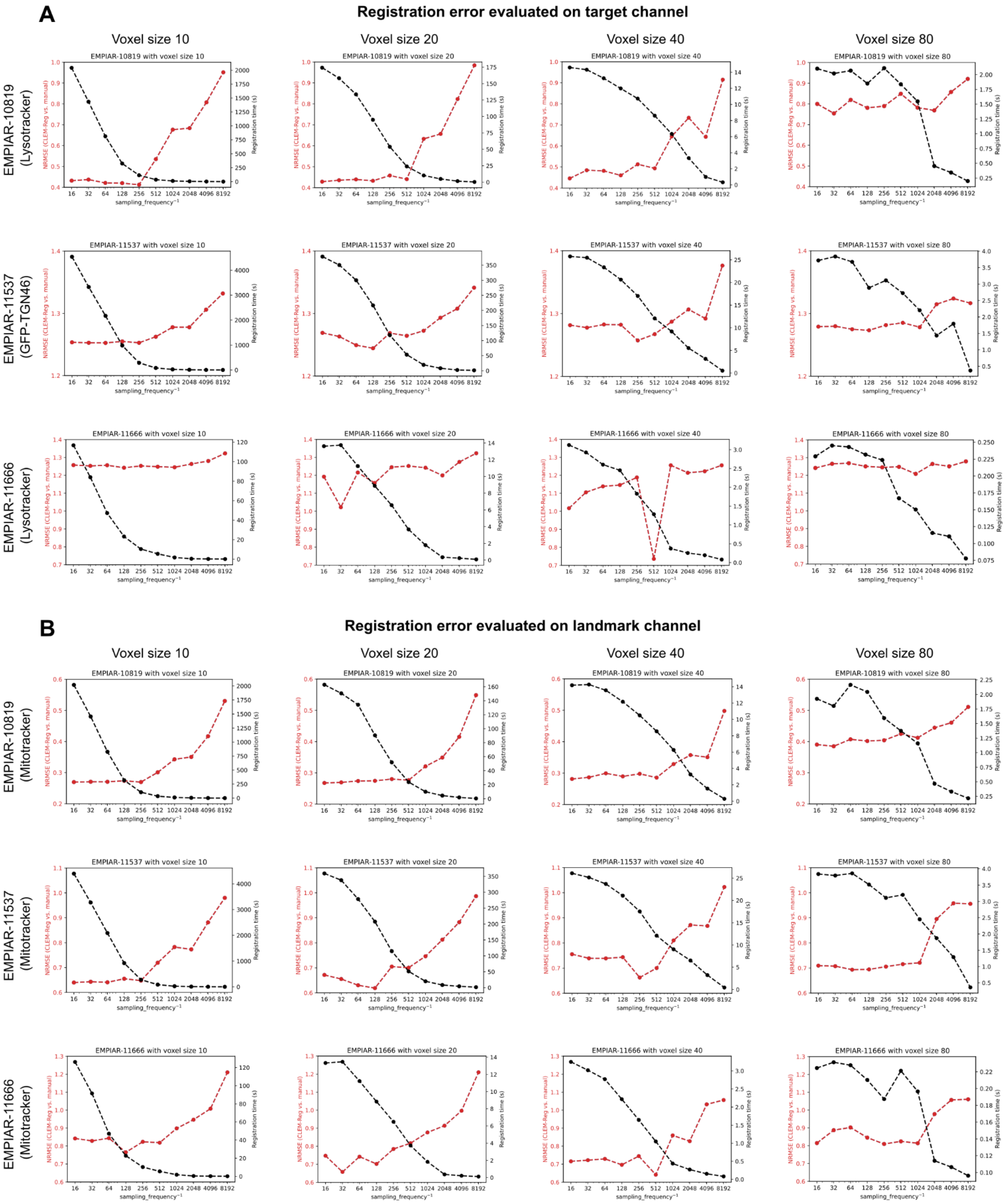


**Supplementary Figure 1. Benchmarking registration performance for varying point cloud sampling parameters.** The normalised root mean squared error (NRMSE) was computed between manually and automatically obtained overlays with CLEM-Reg for a decreasing number of sampled points and increasing voxel sizes across all three benchmark datasets. (A) NRMSE computed on off-target channels (EMPIAR-10819 and EMPIAR-11666: Lysotracker; EMPIAR-11537: GFP-TGN46) and registration time. (B) NRMSE computed on target channel (Mitotracker).

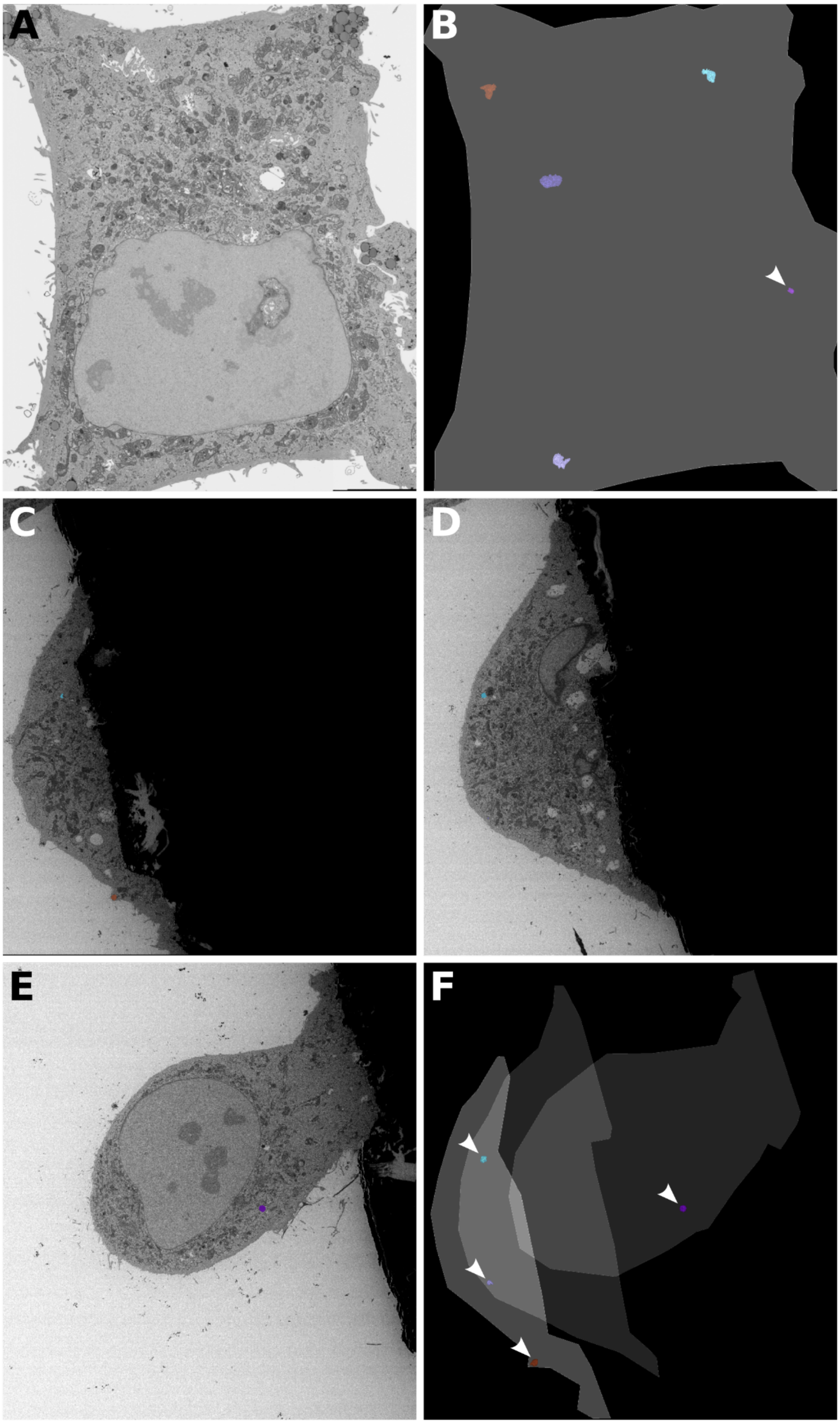


**Supplementary Figure 2. Lysosome distribution across EM volumes**. (A) Representative FIB-SEM slice from EMPIAR-10819 dataset. (B) 3D rendering of cell outline and five manually segmented lysosomes showing distribution across EM volume. Arrowhead points to a small lysosome. (C-E) SBF-SEM slices from EMPIAR-11666 at different z positions. (F) 3D rendering of cell outlines at different z positions and four manually segmented lysosomes showing distribution across EM volume. Arrowheads point to small lysosomes.

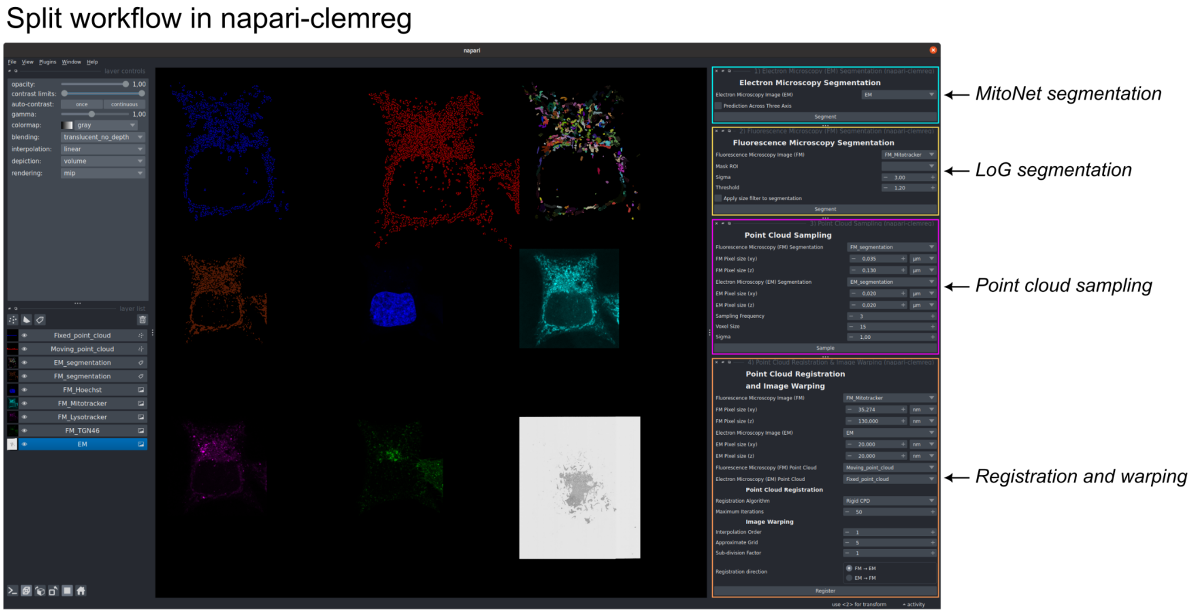


**Supplementary Figure 3. Interface of napari-clemreg split workflow.** Each step of the CLEM-Reg algorithm can be separately executed in the napari-clemreg plugin using the split workflow. This enables users to troubleshoot individual steps, inject intermediate results from external sources and interactively tune individual parameters without executing the full pipeline. Specifically, EM and FM segmentation parameters can be tested, point cloud sampling adjusted and registration and warping settings tested.

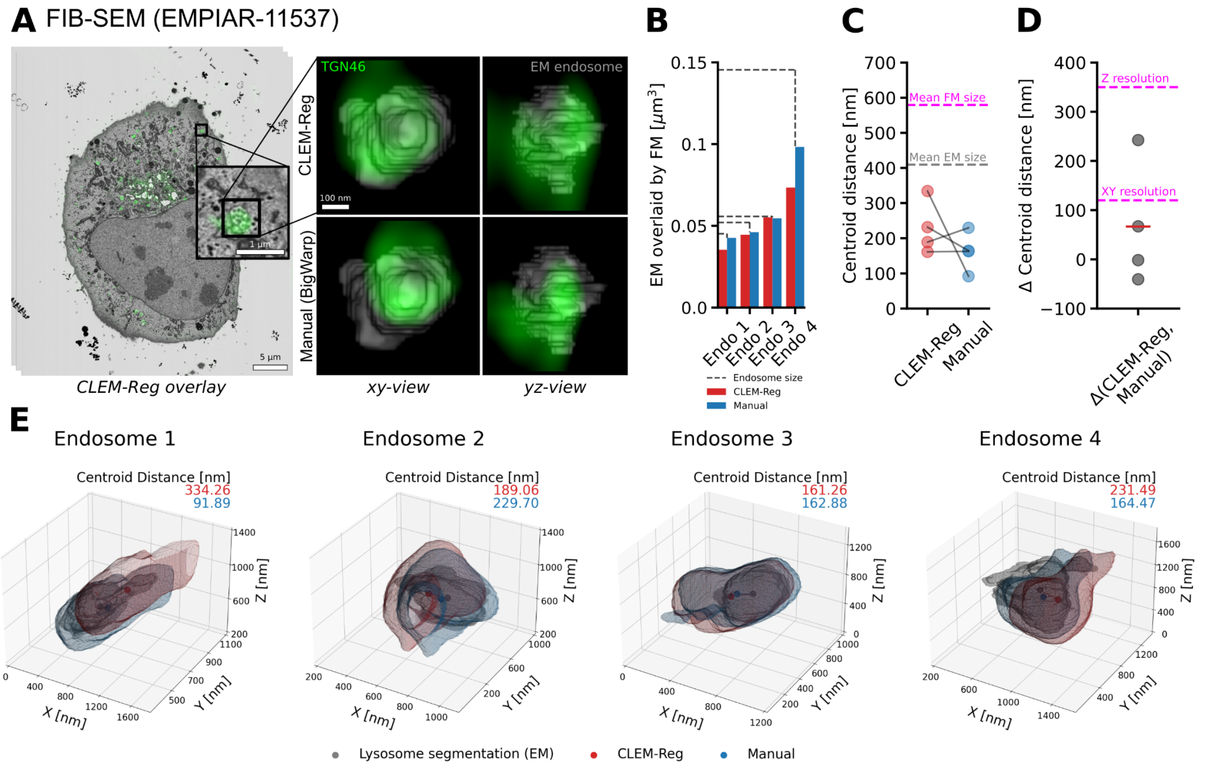


**Supplementary Figure 4. Comparing alignment results between CLEM-Reg and experts**. (A) The GFP-TGN46 channel was overlaid to FIB-SEM (EMPIAR-11537) data using mitochondria as off-target landmarks with CLEM-Reg. To quantify registration performance, four endosomes were manually segmented throughout the EM volume. Corresponding segmentations in FM were obtained by segmenting the GFP-TGN46 channel with Otsu's method. (B) Volume of endosomes in EM overlaid by FM signal was computed by intersecting EM and FM segmentations. (C) Centroid distances between EM segmentations and segmented GFP-TGN46 signal in FM were computed with Euclidean distance. Mean size of FM and EM segmentations are shown in magenta and grey respectively. (D) The difference between GFP-TGN46 signal overlaid manually and with CLEM-Reg was computed from previously found centroid distances. Mean difference in centroid distances is shown with a red horizontal line. The theoretical XY and Z resolution of the fluorescence microscope used is shown in magenta. (E) 3D visualisations of endosome overlays were generated by obtaining meshes from segmentations of EM shown in grey, Lysotracker signal registered using BigWarp (Manual) shown in blue and CLEM-Reg shown in red. Visualisations generated with napari and matplotlib.

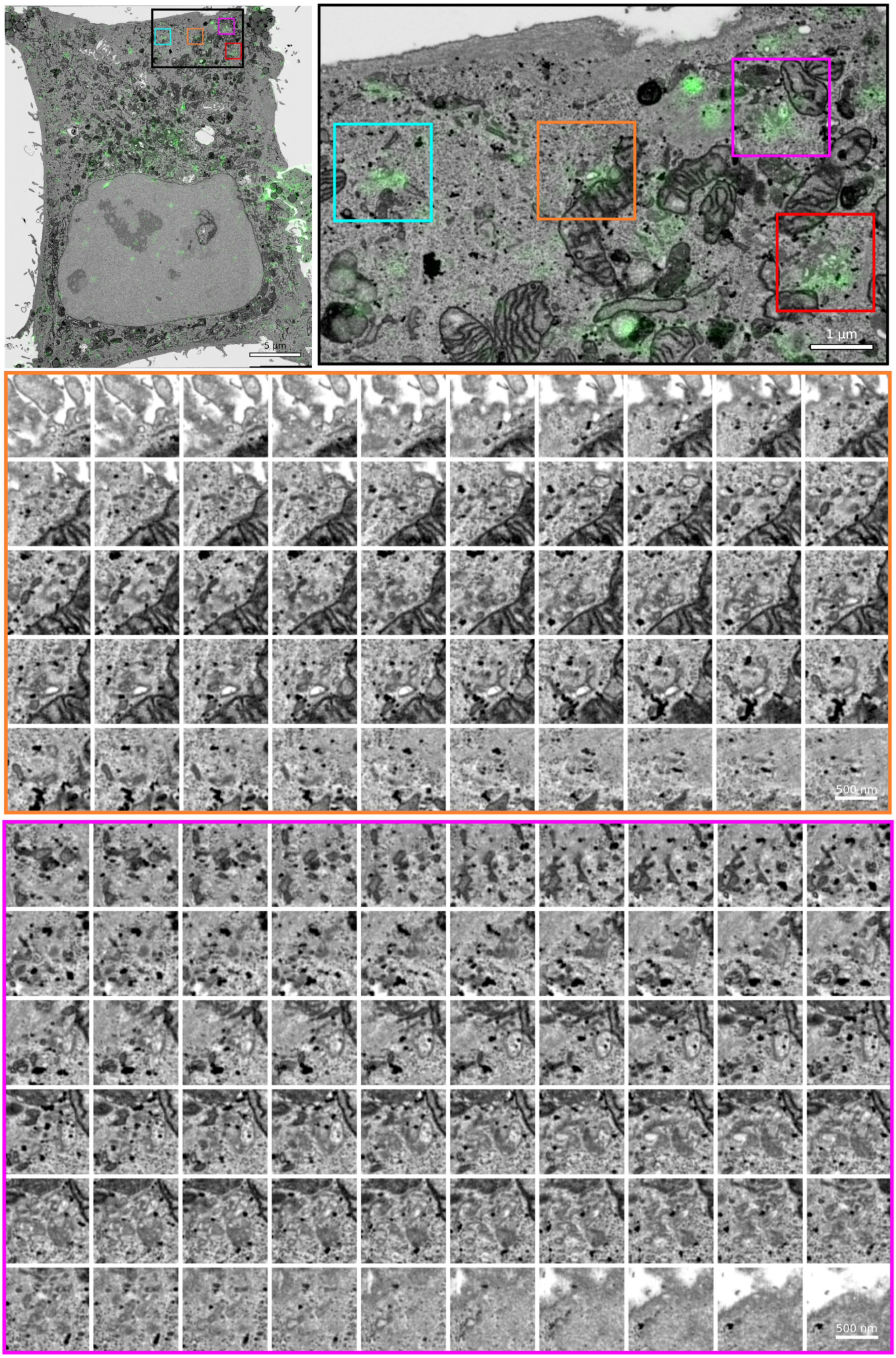


**Supplementary Figure 5. Montage of GFP-TGN-46-positive structures in EMPIAR-10819**. Full montages of GFP-TGN46-positive structures with 10 nm z spacing on EMPIAR-10819. Movies of GFP-TGN46-positive structures shown in **Supp. Movie 1** (orange) and **Supp. Movie 2** (magenta).

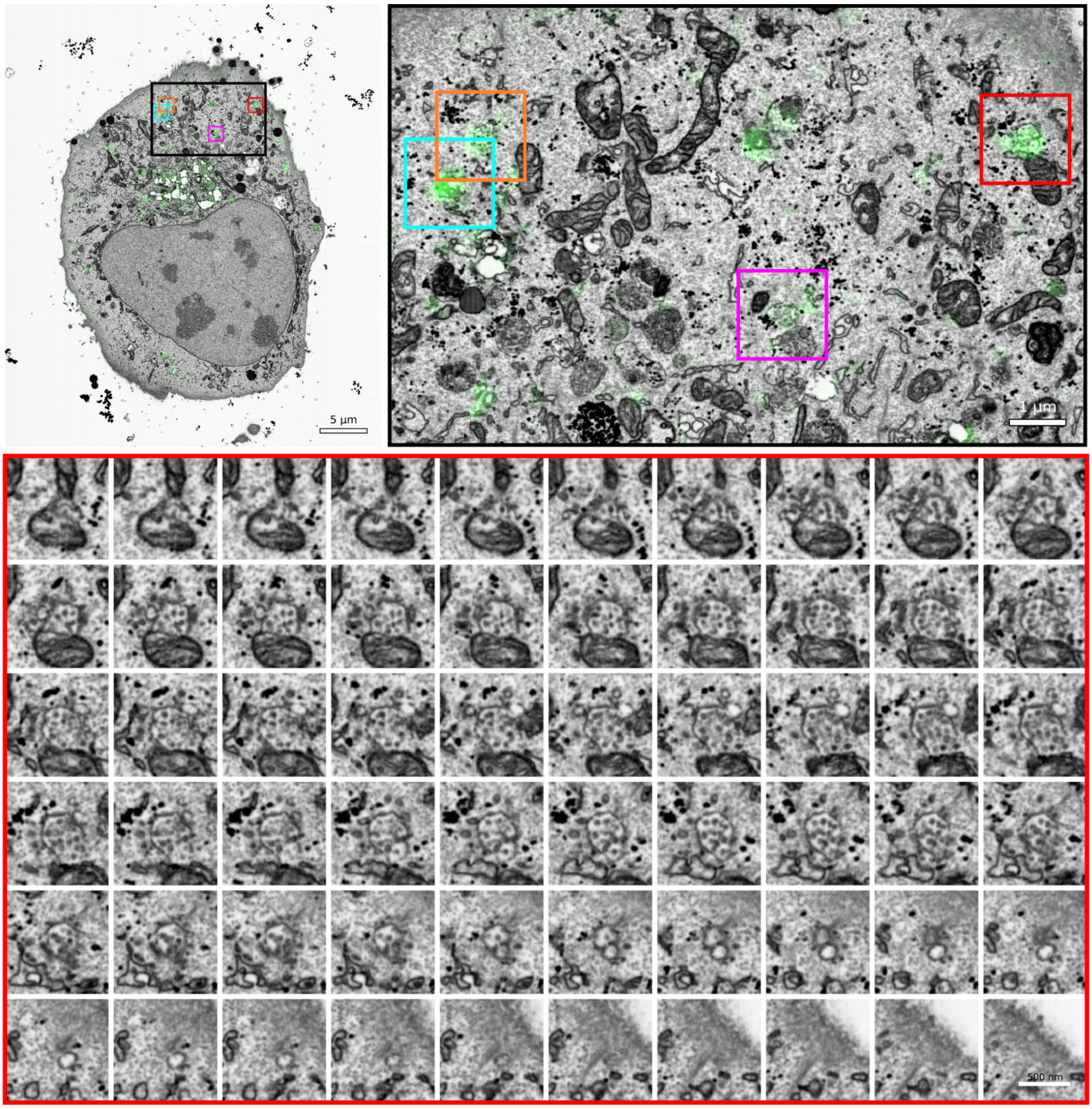


**Supplementary Figure 6. Montage of GFP-TGN-46-positive structures in EMPIAR-11537**. Full montages of GFP-TGN46-positive structures with 10 nm z spacing on EMPIAR-11537. Movie of GFP-TGN46-positive structure shown in **Supp. Movie 3**.

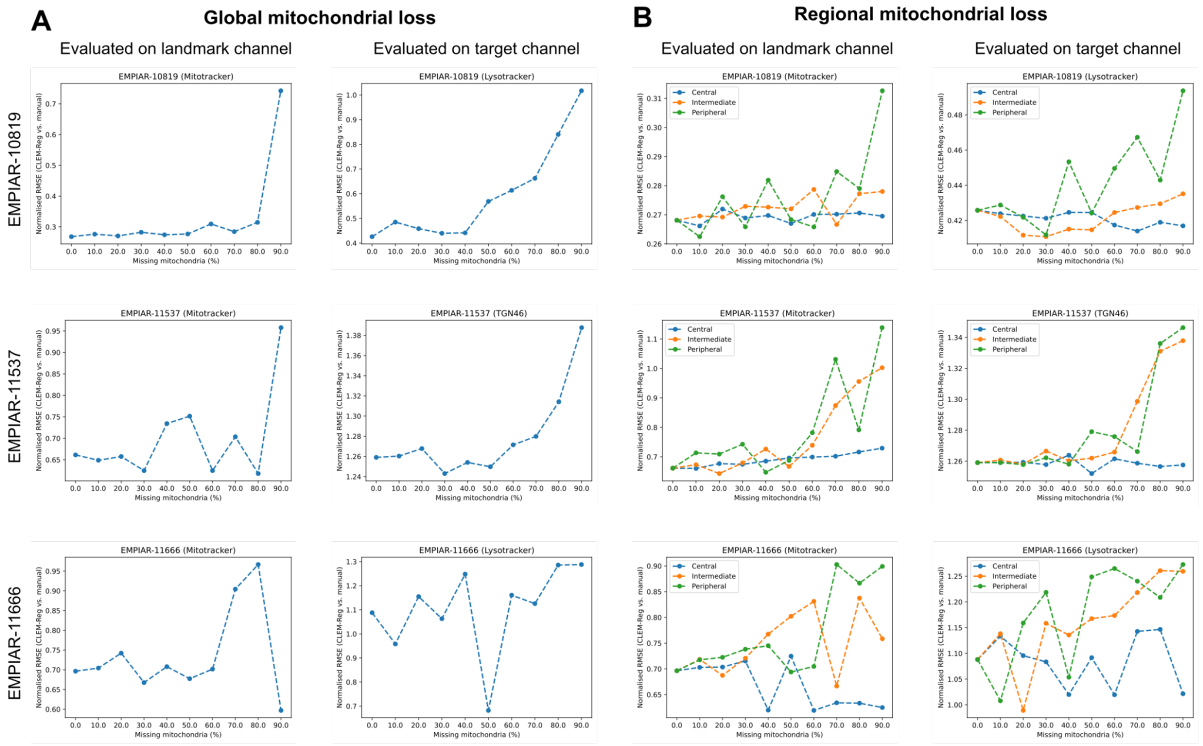


**Supplementary Figure 7. Benchmarking registration robustness to missing mitochondria.** The normalised root mean squared error (NRMSE) was computed between manually and automatically obtained overlays with CLEM-Reg as a function of the percent of missing mitochondria in the EM segmentation mask. A fixed voxel size of 10 and sampling frequency of 1/128 was kept for the point cloud generation. (A) Mitochondria were randomly removed by identifying mitochondrial instances with connected components and then removing mitochondria with the probability indicated on the x-axis. (B) Mitochondria were removed with a probability indicated on the x-axis in three areas with increasing distance from the centre of the image volume (“central” in blue, “intermediate” in orange and “peripheral” in green).
