## Supplementary figures and images for "CLEM-Reg: An automated point cloud based registration algorithm for correlative light and volume electron microscopy"

### Supplementary Movie 1

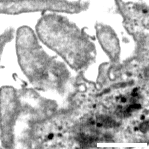

### Supplementary Movie 2

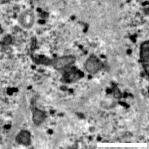

### Supplementary Movie 3

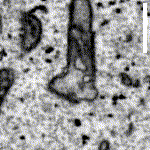
